# Reveel: large-scale population genotyping using low-coverage sequencing data

**DOI:** 10.1101/011882

**Authors:** Lin Huang, Bo Wang, Ruitang Chen, Sivan Bercovici, Serafim Batzoglou

## Abstract

Population low-coverage whole-genome sequencing is rapidly emerging as a prominent approach for discovering genomic variation and genotyping a cohort. This approach combines substantially lower cost than full-coverage sequencing with whole-genome discovery of low-allele-frequency variants, to an extent that is not possible with array genotyping or exome sequencing. However, a challenging computational problem arises when attempting to discover variants and genotype the entire cohort. Variant discovery and genotyping are relatively straightforward on a single individual that has been sequenced at high coverage, because the inference decomposes into the independent genotyping of each genomic position for which a sufficient number of confidently mapped reads are available. However, in cases where low-coverage population data are given, the joint inference requires leveraging the complex linkage disequilibrium patterns in the cohort to compensate for sparse and missing data in each individual. The potentially massive computation time for such inference, as well as the missing data that confound low-frequency allele discovery, need to be overcome for this approach to become practical. Here, we present Reveel, a novel method for single nucleotide variant calling and genotyping of large cohorts that have been sequenced at low coverage. Reveel introduces a novel technique for leveraging linkage disequilibrium that deviates from previous Markov-based models. We evaluate Reveel’s performance through extensive simulations as well as real data from the 1000 Genomes Project, and show that it achieves higher accuracy in low-frequency allele discovery and substantially lower computation cost than previous state-of-the-art methods.

## Introduction

Identification of genomic variation in human DNA sequences is a key first step in associating alleles with human traits and diseases (The 1000 Genomes Project Consortium 2012). Genome-wide Association Studies (GWAS) have successfully linked genetic variation across thousands of genotyped individuals and hundreds of traits (Feero and Guttmacher 2010, Franke et al. 2010, Hindorff et al. 2009, The Wellcome Trust Case Control Consortium 2007). Beyond human, association of genomic variations with traits has many applications, such as in the quality breeding of plants and livestock (Feuillet et al. 2011, The Bovine HapMap Consortium 2009, Huang and Han 2014). Despite their success in linking variation with traits, GWAS performed on genotypes have so far failed to explain a large portion of the heritability of common traits and diseases such as diabetes, schizophrenia and heart disease (Manolio et al. 2009, Visscher et al. 2012, Cirulli and Goldstein 2010, Billings and Florez 2010). Genotype-based GWAS have only examined common single nucleotide polymorphisms (SNPs), and one promising avenue for finding the “missing heritability” is the association of rare variants with common traits (Zuk et al. 2014, Lee et al. 2014, Gibson 2012), also known as the “*common disease rare variant*” hypothesis. Many recent efforts have focused on discovering such rare variants in large cohorts through sequencing rather than genotyping (Tennessen et al. 2012).

Algorithms that call SNPs on a single target genome require the sample to be sequenced at a high coverage (>30×) to confidently differentiate alternate alleles from sequencing errors (Bentley et al. 2008, DePristo et al. 2011, McKenna et al. 2010, Li et al. 2009). Such an approach, however, is expensive when applied to large cohorts. Recently, low-coverage sequencing of large cohorts has been proposed as more cost-efficient and informative than sequencing fewer individuals at high coverage (Li et al. 2011). Many on-going projects have adopted this low-coverage strategy, including the UK10K project (UK10K Consortium), the 1000 Genomes Project (The 1000 Genomes Project Consortium 2010), and the Cohorts for Heart and Aging Research in Genomic Epidemiology (CHARGE) Project (CHARGE Consortium 2009). Each project sequences thousands of individuals at a relatively low coverage. For example, the 1000 Genomes Project sequenced 2,535 whole genomes at depth 4-6×; the CHARGE Project sequenced ~5,000 whole genomes at depth 7×.

To leverage the wealth of genomic data that such large-scale population sequencing projects are providing, computational methods that perform accurate and efficient detection and genotyping of rare SNPs in a population are urgently needed. The corresponding computational problem is considerably more challenging than single-sample genotyping from deep sequencing data: to overcome the noise and missing data inherent in low-coverage sequencing, variant detection and genotyping require the joint estimation of all genotypes of all individuals simultaneously, which needs to leverage the linkage disequilibrium (LD) present in the sequenced cohort. As a result, computation time can become prohibitive and accuracy is harder to achieve for rare alleles.

A number of existing computational methods can be applied to population genotyping. Although not designed for analyzing low-coverage sequencing data, SAMtools (Li et al. 2009), GATK Unified Genotyper (DePristo et al. 2011, McKenna et al. 2010) and Beagle (Browning and Browning 2009) can perform population genotyping (The 1000 Genomes Project Consortium 2012). In particular, applying GATK Unified Genotyper to 62 CEU samples from the 1000 Genomes Project pilot phase collectively, followed by Beagle, leads to reasonably accurate genotyping for common polymorphisms (Nielsen et al. 2012). QCALL (Le and Durbin 2011) employs a dynamic programming algorithm to estimate, for every position of the genome, the posterior probability of presence of an alternate allele in the cohort. The QCALL algorithm then constructs a set of possible ancestral recombination graphs from samples to estimate the SNP posterior probability for each site in each sample from these graphs. The glfMultiples-Thunder pipeline employs a hidden Markov model (HMM) that leverages LD information across a population to genotype likely polymorphic sites, and is currently considered the state of the art for accurate genotyping of populations using sequencing data (Li et al. 2011). In the underlying HMM, each hidden state is a pair of reference haplotypes, which are most closely related to the sample being considered, and observations are genotype likelihoods. To apply this HMM on a sequenced cohort, the sequenced individuals are used as references.

Despite their considerable success, existing genotyping methods are not ideally suited for application to large cohorts (5,000 – 1,000,000 individuals) because of their prohibitive computation time, as well as their reduced accuracy when calling low-frequency genomic variants, which are hard to differentiate from sequencing errors. For instance, the HMM model underlying Thunder links polymorphic sites to surrounding mosaics, modeling these links using a first-order Markovian model. However, the presence of low frequency (0.5% to 5%) and rare (< 0.5% frequency) variants hierarchically breaks the common haplotypes into many uncommon or rare haplotypes, reducing the fit to a model with an underlying Markovian assumption. Additionally, given a cohort of size *n*, the HMM requires *O*(*n*^2^) hidden states, which results in prohibitively high computational overhead as *n* increases. The SNP detection dynamic programming algorithm of QCALL, on the other hand, is more computationally efficient because it does not account for the non-random associations between loci, but its accuracy is reduced for the same reason.

Here, we present Reveel, a novel method for large-scale SNP discovery and genotype imputation using low-coverage sequencing data sets. Reveel leverages the underlying complex LD structure by employing a simplified model that scales linearly with the number of individuals in a cohort for a given number of imputed SNPs, while producing highly accurate genotype calls for both high-and low-frequency SNPs. We evaluate the performance of Reveel on simulated data, as well as real data, and demonstrate that Reveel achieves significant improvements in both efficiency and accuracy over previous state-of-the-art population-scale genotyping methods. We further show that Reveel’s accuracy improves as the cohort size increases, while the computation time scales linearly, making Reveel a practical approach for large-scale population genotyping.

## Results

### Overview of algorithms

Given a cohort of *n* sequenced individuals, with read counts for every allele across the genome of every sample, Reveel genotypes the individuals with a summarization-maximization algorithm that has similarities in form with expectation maximization (EM), through the following five steps: (1) Identification of *m* candidate polymorphic sites across the genome; those are sites where a number of individuals from the cohort may exhibit a minor allele. (2) Initialization of genotypes **G** = (*g*_*i,j*_), where *g*_*i,j*_ is the count {0, 1, 2} of minor alleles that individual *i* has in candidate site *j*. (3) Calculation of a matrix **P** = (*p*_*i,j,g*_), representing the probability of individual *i* having genotype *g* in position *j* given the current assignment **G**. (4) Calculation of new assignment **G’** that maximizes the current entries of **P**; steps 3 and 4 are performed iteratively until convergence. (5) Final refinement of the genotypes **G**.

A key feature of the algorithm is the way in which LD information is leveraged, primarily in step 3, to inform site *j* of individual *i* by taking into account sites *j*’ that are in LD with the variant at location *j* across all individuals. Formally, we initialize a graph in which nodes represent the *m* candidate polymorphic sites and edges represent LD between sites (Figure 1A). For each site *j*, we select *k* sites that exhibit the highest LD with *j*, referred to as *k nearest neighbor* sites (Figure 1B). The criterion for selecting these sites is based on the Jaccard index, namely, intersection over union among the sets *S*_*j*_ and *S*_*j'*_ of individuals in the cohort that exhibit alternate alleles at sites *j* and *j*’, respectively; this computationally efficient approach provides a practical way to select informative sites (see Methods for details). By default, *k* is 2-5 depending on the cohort size *n*. In the rest of the paper, we call *nearest neighbors* of a site *j* the sites that exhibit the strongest LD with *j*, in contrast with the sites physically close to *j*. The corresponding *k* edges are kept in the graph; the remaining edges are removed.

**Figure 1.**
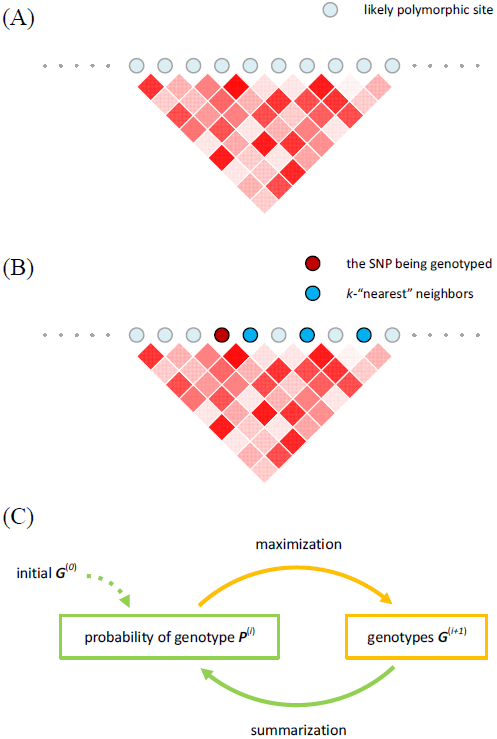
An overview of Reveel. (A) The underlying network of Reveel is composed of a set of likely polymorphic sites and the linkage disequilibrium among them. (B) For every polymorphic site, we pick its *k*-“nearest” neighbor sites in terms of linkage disequilibrium to facilitate genotype calling at the target site. (C) We infer the genotypes using a summarization-maximization iterative method. In every iteration, we apply the summarization step to every SNP in turn and then apply the maximization to every SNP. The summarization step calculates the genotype probabilities using the current estimation of genotypes and observed reads in the context of linkage disequilibrium. The maximization finds the genotypes that maximizing the genotype probabilities obtained in the summarization step. These genotypes are then used to refine the genotype probabilities in the next summarization step. We iterate these two steps until convergence.

A straightforward estimation of LD between every pair of likely polymorphic sites requires *m*^2^ calculations. As LD generally decreases as a function of distance (Reich et al. 2001, Schaffner et al. 2005), to identify nearest neighbors of every site *j* efficiently, we only evaluate neighbors within a window around *j* of default length 500kb-1Mb, as described in Methods.

Reveel utilizes the above LD graph in steps 2 and 3. In step 2, Reveel initializes the genotype calls *g*_*i,j*_ using the summations of read counts in sample i for reference and alternate alleles at site *j* as well as at *j*’s *k* nearest neighbors. In this way, low read coverage at site *j* is partially overcome by informing the genotype assignment using read coverage in *j*’s nearest neighbors. In step 3, similarly, the probabilities *p*_*i,j,g*_ are approximated by leveraging the LD graph as described in Methods; steps 3 and 4 are repeated until convergence (Figure 1C).

### Performance on Simulated Data based on 1KGP

#### Experimental setup

We created a simulated data set, *1kgp-sim*, which mimicked the features of the 1000 Genomes Project (1KGP) dataset (The 1000 Genomes Project Consortium 2010), including high variability of sequencing depth among loci and individuals. *1kgp-sim* included 2,535 samples; each corresponded to a sample in the 1KGP data set. To create these samples, we simulated variants in 10,000 haplotypes for a 1-Mbp region using COSI with parameters from the best-fitting model (Schaffner 2005). A 1-Mbp region on chromosome 20 (43,000,000-44,000,000) of the human genome build GRCh37 was used as a reference genome. Combining these variants with the reference genome resulted in 10,000 chromosomes. A simulated sample was a composition of two randomly selected simulated chromosomes. Then, for each sample, we downloaded the BAM files from the 1KGP database to obtain the mapping position and length of each real read, and we generated a simulated read with the same position and length, and with sequencing base errors injected into a haplotype of the simulated sample; the sequencing base error rate was set to 0.1% (Lou et al 2013), which we estimated from the 1KGP data as explained in Methods. The base qualities of the simulated reads were copied from the downloaded bam files. Finally, the simulated reads were mapped to the reference genome by BWA. The resulting bam files (as opposite to the bam files downloaded from the 1KGP) were used in this set of experiments. We generated three additional data sets, *1kgp-sim-n100*, *1kgp-sim-n500*, and *1kgp-sim-n1000*, from *1kgp-sim* by using randomly selected 100, 500, and 1000 individuals respectively.

#### SNP discovery

First, we measured the ability of Reveel, GATK, and glfMultiples, to identify sites in the genome that are polymorphic in the samples. Reveel showed a near-perfect performance for discovering common SNPs (Supplementary Figure S1). In Figure 2 we illustrate the performance of Reveel in detecting rare and low frequency SNPs, and compare with performance of GATK and glfMultiples. SNPs were divided into three bins according to their allele frequencies (AF): < 0.1%, 0.1 −0.2% and 0.2 −0.5%. For each bin, we report the recall of each method, as the fraction of SNPs that were identified among all SNPs in the bin. We report precision as the fraction of identified loci that showed more than one allele in the simulated 10,000 haplotypes out of all reported loci. To evaluate glfMultiples, we set the minimum and maximum values of the average total depth per individual as 0.5 and 20, following the authors’ recommendation. The posterior cutoff varied from 0.5 to 1 to obtain a ROC curve. We ran GATK with default parameters, except that we varied the minimum PhRED-scaled confidence threshold for variant calling from 15 to 40, to obtain ROC curves.

**Figure 2.**
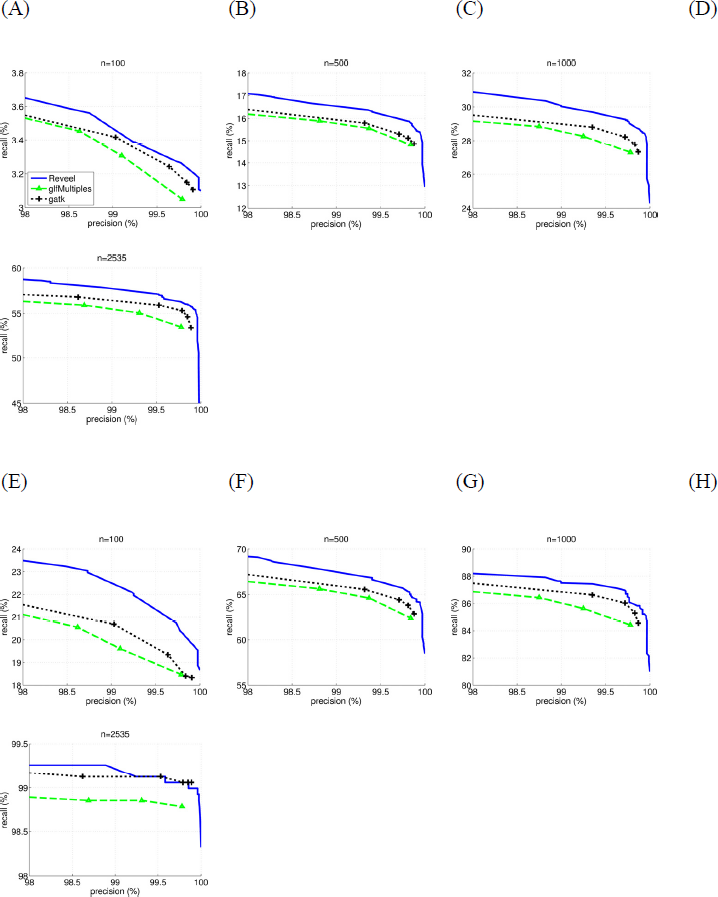
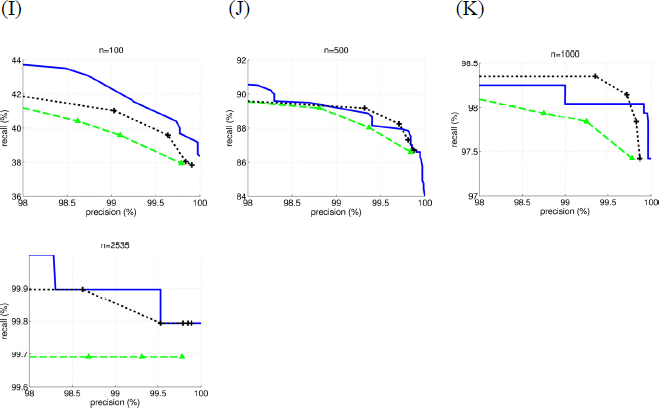
Performance of Reveel in discovering rare and low-frequency SNPs. SNPs were called on the simulation data sets using the following methods: Reveel, glfMultiples (Li et al. 2011), and GATK Unified Genotyper (DePristo et al. 2011, McKenna et al. 2010) applied to all the samples collectively. We compared the recall and precision of these methods for discovering the rare and low-frequency SNPs, which were grouped into three sets according to their AF.

As shown in Figure 2, Reveel outperformed other methods in discovering SNPs with AF < 0.1% and SNPs with AF ranging between 0.1 to 0.2%. On SNPs with AF 0.2-0.5%, Reveel showed similar recall with GATK for the *n* = 500, 1000, and 2535 cases, and higher recall for the *n* = 100 case. In a large cohort, a SNP with AF 0.2-0.5% is very likely to be captured by multiple reads in a few samples; hence, state-of-the-art callers such as GATK are capable of discovering SNPs of moderate AF.

#### Genotyping

##### Genotyping accuracy

We measured the genotyping accuracy of each method, defined as the percentage of inferred genotypes that are correct. Table 1 and Supplementary Figure S2 demonstrate Reveel's genotyping accuracy measured at a 100% precision level. The performance of Thunder was measured with its default parameters; the SNP calling precisions were 98.62%, 98.81%, and 98.75% for the *1kgp-sim-n100*, *1kgp-sim-n500*, and *1kgp-sim-n1000* cases respectively. As the CPU overhead of Thunder was considerably high when Thunder was applied to *1kgp-sim*, as shown in Table 2, we did not report the genotyping accuracies of Thunder for this data set. For direct comparison, Table 1 shows the genotyping accuracy at the polymorphic sites detected by all three genotype-callers. The genotyping accuracies at sites detected by each caller individually are shown in Supplementary Figure S2. The genotyping accuracy of Reveel approached perfect performance when the sample size increased to thousands. In our experiments, Reveel achieved 99.98% genotyping accuracy on the *1kgp-sim* data set, which included all 2,535 samples. This implies that Reveel will become increasingly powerful as cohort sizes increase in the future.

**Table 1.**
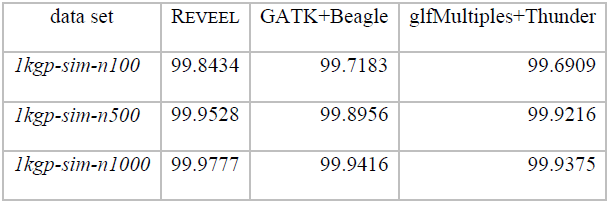
Genotyping accuracy. We evaluated the genotype calling performance of Reveel and two state-of-the-art methods (glfMultiples+Thunder and GATK+Beagle). As these callers reported different SNP sites, for fair comparison we measured the genotyping accuracies at the SNPs discovered by all three callers.

**Table 2.**
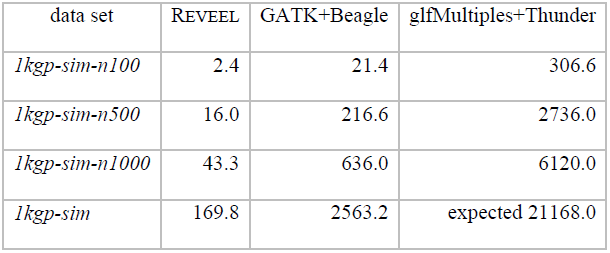
Running time. This table lists the CPU overhead, measured in minutes, of both SNP discovery and genotype calling; the genotyping step represents the major bottleneck.

##### Computation time

We compared the running time of Reveel to other methods on a 2.67GHz Intel Xeon X5550 processor, as shown in Table 2. Our method was more than 120 times faster than glfMultiples+Thunder, and 9 to 15 times faster than GATK + Beagle. The process of finding *k*-nearest neighbors for *m* polymorphic sites has a time complexity of *O*(*nm*^2^), which we further reduce by restricting the calculation to blocks of size <1Mbp; the time complexity of the iterative algorithm is *O*(*nm*).

##### Performance on uncommon SNPs

We grouped SNPs according to their AFs and compared the performance of tools on each group (Figure 3). Reveel exhibits higher accuracy than GATK+Beagle and glfMultiples+Thunder. Because the reported genotyping accuracy was dominated by the large number of homozygous reference sites, we also measured the performance of tools on calling alternate alleles on each group (Table 3). Again, Reveel exhibited higher accuracy than the other two methods on most cases. We also grouped SNPs in *1kgp-sim-n1000* according to whether they are homozygous reference (hom-ref), heterozygous (het), or homozygous alternate (hom-alt), and reported accuracy in each class (Table 4). It has been previously suggested that high genotyping accuracy at sites with low AF can be achieved by simply assigning homozygous reference (also known as a “straw-man” approach) (Li et al. 2011). Table 4 shows that Reveel rarely calls alternate alleles as reference.

**Figure 3.**
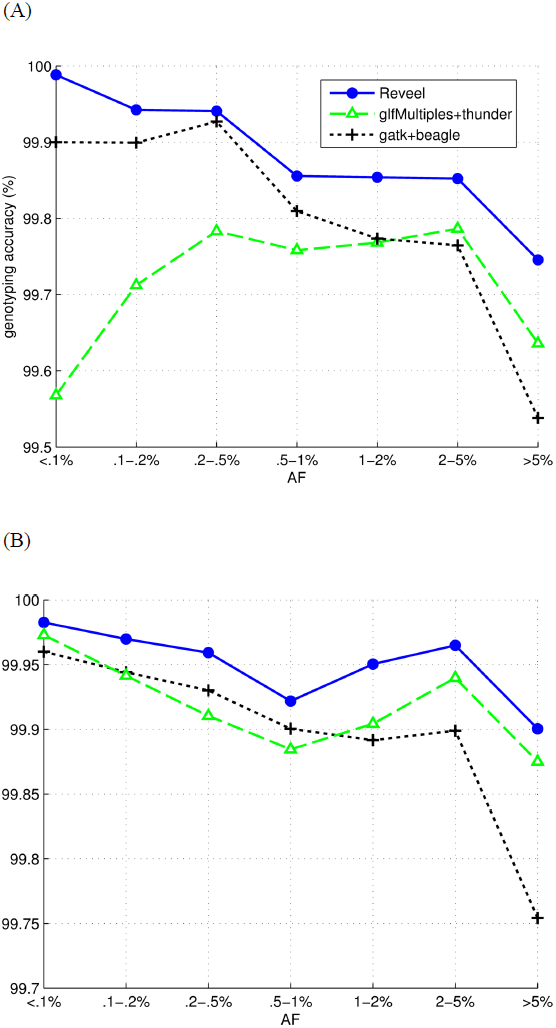
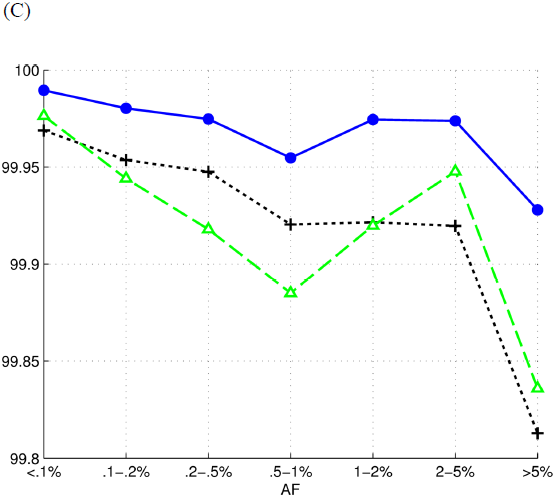
Genotyping performance as a function of allele frequencies. The polymorphic sites were categorized according to their population minor allele frequencies, which were computed as the percentage of minor alleles in 10,000 simulated haplotypes. We compared the performance of Reveel, glfMultiples followed by Thunder (Li et al. 2011), and GATK Unified Genotyper (DePristo et al. 2011, McKenna et al. 2010) applied to all the samples collectively followed by Beagle (Browning and Browning 2009) at the sites in each category.

**Table 3.** Performance of Reveel in calling alternate alleles. We grouped the polymorphic sites according to their allele frequencies. To measure the efficiency of each group, we reported sensitivity as the percentage of alternate alleles that are correctly identified and precision as the percentage of true alternate alleles against all the called alternate alleles.

| AF<br>category | REVEEL |  | GATK+Beagle |  | glfMultiples+Thunder |  |
| --- | --- | --- | --- | --- | --- | --- |
|  | recall<br>(%) | precision<br>(%) | recall<br>(%) | precision<br>(%) | recall<br>(%) | precision<br>(%) |
| <.1% | 99.1304 | 99.7812 | 99.6785 | 97.9463 | 99.8945 | 90.2334 |
| .1-.2% | 96.9697 | 99.5556 | 99.0836 | 97.6298 | 99.7753 | 91.1704 |
| .2-.5% | 97.7337 | 99.1379 | 99.1701 | 97.0230 | 99.8639 | 89.7311 |
| .5-1% | 97.1652 | 98.8072 | 97.2831 | 97.7974 | 99.1304 | 94.7631 |
| 1-2% | 97.5171 | 99.6065 | 97.5960 | 97.8436 | 99.4125 | 96.0276 |
| 2-5% | 98.8254 | 99.6638 | 98.6744 | 98.9367 | 99.6416 | 98.2181 |
| $\geq 5\%$ | 99.6754 | 99.7380 | 99.4361 | 99.4973 | 99.7031 | 99.4581 |

(B) $n = 500$
| AF<br>category | REVEEL |  | GATK+Beagle |  | glfMultiples+Thunder |  |
| --- | --- | --- | --- | --- | --- | --- |
|  | recall<br>(%) | precision<br>(%) | recall<br>(%) | precision<br>(%) | recall<br>(%) | precision<br>(%) |
| <.1% | 99.4073 | 97.3668 | 99.3221 | 96.3671 | 99.1008 | 97.8567 |
| .1-.2% | 97.9528 | 98.5997 | 97.6410 | 96.4064 | 96.2209 | 97.3529 |
| .2-.5% | 96.7025 | 99.0314 | 96.5767 | 95.9772 | 94.4334 | 95.8870 |
| .5-1% | 97.5731 | 99.4949 | 98.0930 | 98.1414 | 97.4372 | 98.1791 |
| 1-2% | 99.1637 | 99.7436 | 98.8762 | 98.7346 | 99.4186 | 98.4851 |
| 2-5% | 99.7722 | 99.8999 | 99.5498 | 99.5028 | 99.9083 | 99.5295 |
| $\geq 5\%$ | 99.8768 | 99.8944 | 99.6818 | 99.7534 | 99.9043 | 99.8087 |

(C) $n = 1000$
| AF<br>category | REVEEL |  | GATK+Beagle |  | glfMultiples+Thunder |  |
| --- | --- | --- | --- | --- | --- | --- |
|  | recall<br>(%) | precision<br>(%) | recall<br>(%) | precision<br>(%) | recall<br>(%) | precision<br>(%) |
| <.1% | 98.9421 | 98.1873 | 98.8715 | 95.1493 | 96.8247 | 94.6213 |
| .1-.2% | 97.9689 | 99.3920 | 97.4858 | 96.0681 | 92.7237 | 95.4523 |
| .2-.5% | 97.6187 | 99.3227 | 97.2031 | 96.4517 | 92.6783 | 95.7634 |
| .5-1% | 98.6654 | 99.6529 | 98.4520 | 98.5936 | 95.9814 | 96.9489 |
| 1-2% | 99.5954 | 99.8394 | 99.2068 | 99.0520 | 99.2445 | 98.1897 |
| 2-5% | 99.8507 | 99.9068 | 99.6383 | 99.6192 | 99.7875 | 99.3575 |
| $\geq 5\%$ | 99.9082 | 99.9261 | 99.7635 | 99.8062 | 99.6830 | 99.5582 |

**Table 4.** Genotype error rates. This table shows the average genotype error rates when applying Reveel to the *1kgp-sim-n1000* data set.

| truth | outcome |  |  | error rate |
| --- | --- | --- | --- | --- |
|  | hom-ref | het | hom-alt |  |
| hom-ref | 8,131,407 | 873 | 0 | 0.011% |
| het | 1,292 | 652,152 | 208 | 0.229% |
| hom-alt | 1 | 330 | 301,737 | 0.110% |

##### Performance as a function of sequencing error rate.

Reveel requires the sequencing error rate as an input parameter; however, its performance is robust even when the true error rate is different from the one given as input: we created three simulated data sets in which the injected sequencing base error rates were 0.05%, 0.1%, and 0.2% respectively. The input sequencing error rate of Reveel was set to be 0.1% for all three data sets. The performance of the 0.05% and 0.2% cases remains on the same level as the 0.1% case, as shown in Supplementary Table S1.

### Performance on 1KGP samples

We applied Reveel to the low-coverage sequencing data from the 1000 Genomes Project. This data set includes 2,535 samples from 26 populations (Supplementary Table S2). We restricted our analysis to a 5-Mbp region on chromosome 20 (43,000,000-48,000,000), which we call *1kgp-real*. Reveel was applied to call SNPs and genotypes from each population separately. The block size was set to 500-kb. As a post-processing step, we merged SNPs detected in each population, and we reported genotypes of each sample on the merged SNP set 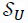. The genotype of a sample from population *p* at a SNP that belongs to 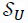 but not 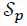 was treated as a homozygous major within population *p*.

#### SNP discovery

Reveel discovered 171,734 likely polymorphic sites in all 26 populations, including 64,724 SNPs reported in the 1KGP Phase 1 and 107,010 putative SNPs. The 1KGP Phase 1 reported 68,208 SNPs in the analyzed region; our method confirmed 94.89% of those. The putative SNPs were primarily rare variants (Figure 4): more than 95% of the putative SNPs were with allele frequencies ⩽ 1%; only 1.5% of putative SNPs had allele frequencies > 5%. Interestingly, Reveel discovered 1,989 triallelic sites and 17 sites having all four nucleotides. The African populations contributed 40,143 putative SNPs, while the other populations showed lower diversity (Table 5).

**Figure 4.**
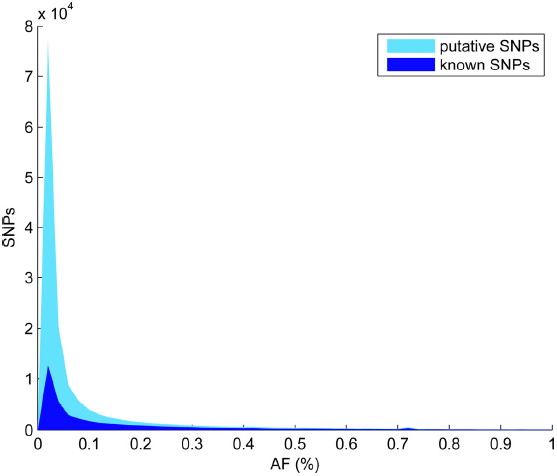
SNPs discovered from 2,535 samples of the 1000 Genomes Project. We plotted the histogram of the allele frequencies of the SNPs discovered by Reveel from 2,535 1KGP samples. The vast majority of common SNPs were reported in the 1000 Genomes Project Phase 1. The putative SNPs are primarily rare ones with AF ≤ 1%.

**Table 5.** The SNPs discovered in populations. We applied Reveel to each population from the 1000 Genomes Project separately, and then collected the SNPs discovered from the populations with the same ancestry. Our tool revealed a large number of putative SNPs from the South Asian populations and the African populations.

| population<br>ancestry | number of<br>populations | discovered<br>SNPs | SNPs reported in<br>Phase 1 | putative<br>SNPs |
| --- | --- | --- | --- | --- |
| East Asian | 5 | 48,090 | 24,144 | 23,946 |
| South Asian | 5 | 55,321 | 20,653 | 34,668 |
| African | 7 | 83,786 | 43,643 | 40,143 |
| European | 5 | 48,406 | 27,293 | 21,113 |
| Americas | 4 | 50,821 | 33,944 | 16,877 |

We compared the SNPs discovered by Reveel and glfMultiples from 99 CEU samples and 109 YRI samples of the *1kgp-real* data set. The SNPs detected by only one method were compared to the SNPs reported in the CEU and YRI trios from the 1000 Genomes Project Pilot 2, because these trios are sequenced at high depth (42x on average) and their genotype calls are likely to be of high accuracy (The 1000 Genomes Project Consortium 2010, Xu et al 2012) and consequently any putative SNPs detected by either method that are also called in these trios have strong evidence of being true. As shown in Table 6, both methods discovered 22,828 SNPs in the CEU trio and 36,527 SNPs in the YRI trio. In addition, each method identified a number of SNPs not found by the other method. Of those, glfMultiples identified more than twice as many as Reveel. The vast majority (~99%) of SNPs identified by only one method were not identified by the deep trio sequencing, and that proportion was slightly higher for glfMultiples, which is consistent with Reveel having a lower false positive rate.

**Table 6.**
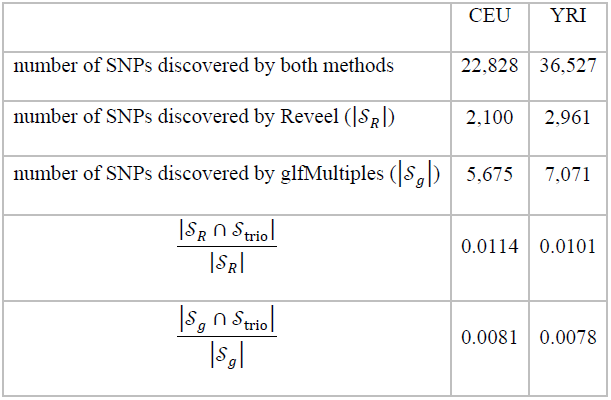
Putative SNPs discovered from the 1kgp-real data set. We compared the SNPs discovered by Reveel and glfMultiples from the CEU and YRI samples of the *1kgp-real* data set. Sequentially, we examined if the SNPs discovered by only one method were detected from the CEU trio and the YRI trio in the 1000 Genomes Project Pilot 2.

|  | CEU | YRI |
| --- | --- | --- |
| number of SNPs discovered by both methods | 22,828 | 36,527 |
| number of SNPs discovered by Reveel ($\vert \mathcal{S}_R\vert $) | 2,100 | 2,961 |
| number of SNPs discovered by glfMultiples ($\vert \mathcal{S}_g\vert $) | 5,675 | 7,071 |
| $\frac{\vert \mathcal{S}_R \cap \mathcal{S}_{\text{trio}}\vert }{\vert \mathcal{S}_R\vert }$ | 0.0114 | 0.0101 |
| $\frac{\vert \mathcal{S}_g \cap \mathcal{S}_{\text{trio}}\vert }{\vert \mathcal{S}_g\vert }$ | 0.0081 | 0.0078 |

#### Genetic diversity among populations

There is a rich literature on the estimation and interpretation of *F*_*ST*_ (Wright 1949, Nei 1973, Holsinger and Weir 2009, Xu et al 2009). We measured the genetic divergence between two populations by using Hudson’s estimator of *F*_*ST*_ (Hudson et al. 1992) interpreted in (Bhatia et al. 2013) over SNPs that Reveel ascertained as polymorphic in any of these 26 populations. Following the strategy used in HapMap 3 (Altshuler et al. 2010) and recommended previously (Weir and Cockerham 1984, Bhatia et al. 2013), we combined the estimates of *F*_*ST*_ across multiple SNPs as the ratio of averaged numerator and averaged denominator of the single SNP *F*_*ST*_ estimates. A single SNP *F*_*ST*_ estimate between populations *A* and *B* is given by

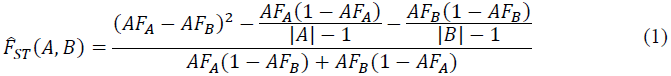

where |*A*| is the sample size of population *A*. The resulting genetic divergence between every pair of populations is shown in Figure 5A. We observed a strong similarity between the populations from the same population ancestry. For example, both ASW (African Ancestry in Southwest US) and ACB (African Caribbean in Barbados) exhibited high similarity with YRI (Yoruba in Ibadan, Nigeria). The two dimensional histograms in Figure 5 compare allele frequencies between pairs of populations. As another example, CDX (Chinese Dai in Xishuangbanna, China) was very similar to KHV (Kinh in Ho Chi Minh City, Vietnam) in terms of the Euclidian distance of allele frequencies. Interestingly, their similarity was stronger than CDX and CHB (Han Chinese in Bejing, China). All the populations with South Asian Ancestry, as expected, exhibited high similarity. Populations with Americas Ancestry, however, exhibited relatively higher divergence, except for CLM (Colombian in Medellin, Colombia) and PUR (Puerto Rican in Puerto Rico). PUR exhibited a relatively high similarity with a few European populations, including IBS (Iberian populations in Spain), CEU (Utah Residents (CEPH) with Northern and Western European ancestry), FIN (Finnish in Finland), GBR (British in England and Scotland), providing a hint to the demographics. As can be seen in Fig. 5(I), at the sites where PEL (Peruvians from Lima, Peru) had low allele frequencies (< 5%), MSL (Mende in Sierra Leone) can have much higher allele frequencies. Furthermore, PEL showed a considerable divergence from all other populations except for MXL (Mexican Ancestry in Los Angeles, California).

**Figure 5.**
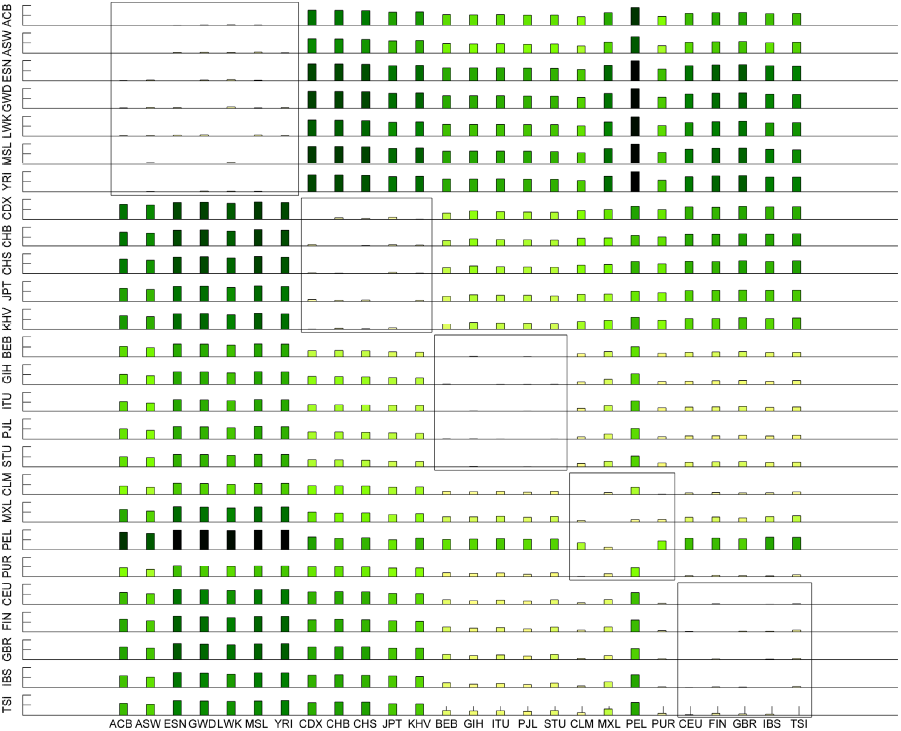
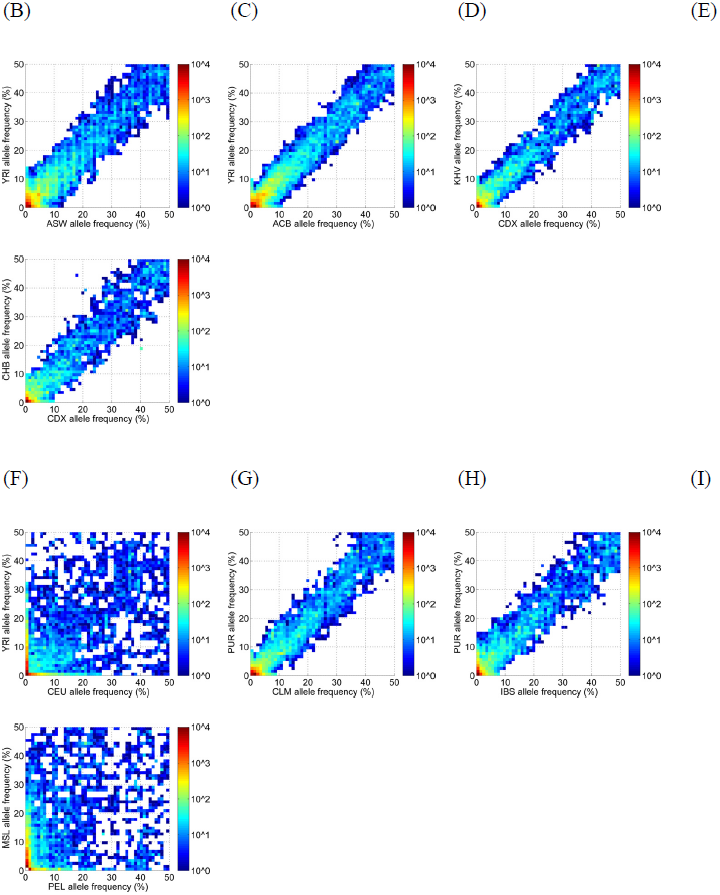
Genetic diversity among populations. (A) The genetic diversity *F*_*ST*_ between every pair of populations, measured over all SNPs ascertained by Reveel as polymorphic in the *1kgp-real* data set. PEL and YRI exhibited the highest *F*_*ST*_ 0.1608 among all the pairs. (B)-(I) Two-dimensional histograms illustrate the comparison of allele frequencies in eight sample pairs of populations.

#### Genotyping accuracy

We evaluated performance on the genotype calls using the genotypes reported in the HapMap Phase III panel (Altshuler et al. 2010) as benchmarks. Out of 26 populations in the *1kgp-real* data set, HapMap 3 studied nine populations: ASW, CEU, CHB, GIH, JPT, LWK, MXL, TSI, and YRI. The number of the common samples between HapMap 3 and 1KGP in these populations were 50, 90, 94, 93, 97, 90, 56, 96, and 103 respectively. Reveel achieved high accuracy on two European populations and three Asian populations (Figure 6). Performance on the ASW and MXL was lowest, perhaps due to the fact that there are only 66 and 67 samples for these populations, respectively.

For comparison, we applied GATK+Beagle and glfMultiples+Thunder to the same data set *1kgp-real*. Similar to the application of Reveel, we merged the SNP sets called from all 26 populations and evaluated the genotyping accuracy at the union SNP set using the HapMap 3 benchmarks. Whenever no tool reported a locus as a SNP, we assumed all the samples were homozygous reference at this locus, where the reference allele came from the reference genome. Figure 6 shows that Thunder and Reveel perform similarly, whereas in simulations Reveel outperforms Thunder. One possible explanation for the discrepancy is that the SNPs reported by HapMap 3 are primarily common SNPs (Table 7), in which the two methods have similar performance in simulations.

**Figure 6.**
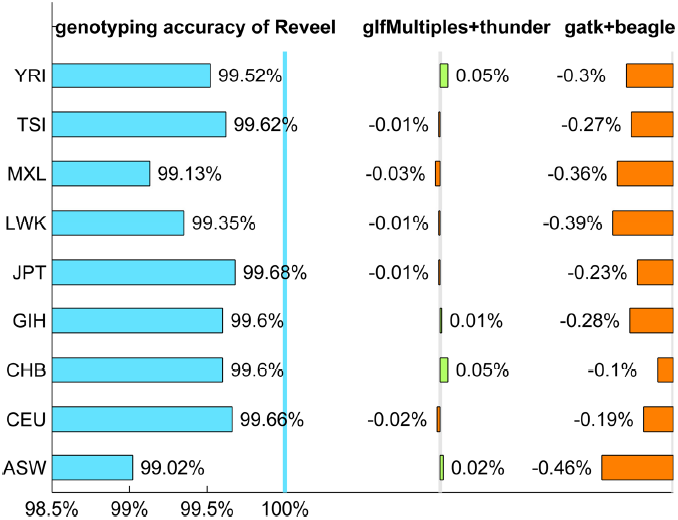
Genotyping accuracy. Genotyping accuracy was evaluated using the genotypes of 50 ASW samples, 90 CEU samples, 94 CHB samples, 93 GIH samples, 97 JPT samples, 90 LWK samples, 56 MXL samples, 96 TSI samples, and 103 YRI samples reported by HapMap 3 as the benchmark.

**Table 7.** Allele frequencies of benchmark SNPs in 1KGP data. We called genotypes from a 5-Mbp region using Reveel, glfMultiples+Thunder, and GATK+Beagle. Out of the polymorphic sites reported by each of these tools, 3,086 sites were also claimed to be SNPs by HapMap3. This table shows the allele frequencies of these 3,086 sites.

| AF category | number of SNPs |
| --- | --- |
| <.1% | 2 |
| .1-.2% | 6 |
| .2-.5% | 8 |
| .5-1% | 16 |
| 1-2% | 84 |
| 2-5% | 284 |
| ≥ 5% | 2686 |
| sum | 3086 |

#### Discordance of alleles between HapMap3 and 1KGP

We also observed a few sites where alleles between HapMap3 and 1KGP are discordant (Table 8). Four sites where alleles called by HapMap3 and 1KGP (type 1) were discovered by both Reveel and GATK+Beagle, and matched the report of a previous publication (Qin et al. 2013). Three alleles where the allele frequencies are considerably different between the genotypes reported by HapMap3 and the genotypes from 1KGP (type 2) were also found by both Reveel and GATK+Beagle, and these were not reported (Qin et al. 2013). For example, at locus chr20: 44697887 HapMap3 reported the vast majority of haplotypes having G (99.53%) and only a small portion having T (0.47%), while Reveel inferred 2.19% G and 97.81% T from 1KGP. At locus chr20:47590564 HapMap3 reported 99.82% C and 0.18% T, while 1KGP exhibited 9.25% C and 90.75% T. At chr20:48661748 although both data sets supported the major allele being A and the minor allele being G, the minor allele frequency reported by HapMap3 was 44.99% and that obtained from 1KGP was only 9.01%. Finally, three loci that were reported as SNPs in HapMap3 and not in 1KGP (type 3), were also reported as constant by both GATK+Beagle and Reveel. When we evaluated the genotyping accuracy of tools, we excluded all the above loci.

**Table 8.** Discordance of alleles between HapMap3 and 1KGP. In the studied 5-Mbp region on chromosome 20 (43,000,000-48,000,000) ten sites exhibited significant discordance in alleles between HapMap3 and 1KGP. We called genotypes at these sites from the 1kgp-real data set using Reveel and GATK+Beagle. The resulting genotypes were compared with the genotypes reported by HapMap3.

| type | locus | alleles in HapMap3 | alleles in 1KGP |
| --- | --- | --- | --- |
| 1 | chr20:43143667 | C/T | A/G |
|  | chr20:43536455 | A/G | C/T |
|  | chr20:44688665 | C/T | A/G |
|  | chr20:46431392 | A/C | G/T |
| 2 | chr20:44697887 | G/T | T/G |
|  | chr20:47590564 | C/T | T/C |
|  | chr20:48661748 | A/G | A/G |
| 3 | chr20:43104944 | C/T | G |
|  | chr20:43850800 | G/A | C |
|  | chr20:44272298 | A/C | T |

## Discussion

A rare genetic variant that originated from a recent mutation event tags many of the other genetic variants surrounding it, as these were present at that time, including variants at long genetic distance from it; rare variants present an extremely high LD, yielding long rare haplotypes. The nearest-neighbor concept in Reveel uniquely leverages this observation: common SNPs tend to have nearest neighbors that are proximal in genetic distance, while rare SNPs tend to have nearest neighbors that are much more distant (Figure 7, Figure 8A); moreover, the allele frequencies of target SNPs and their nearest neighbors are in almost perfect linear correlation (Figure 8B).

**Figure 7.**
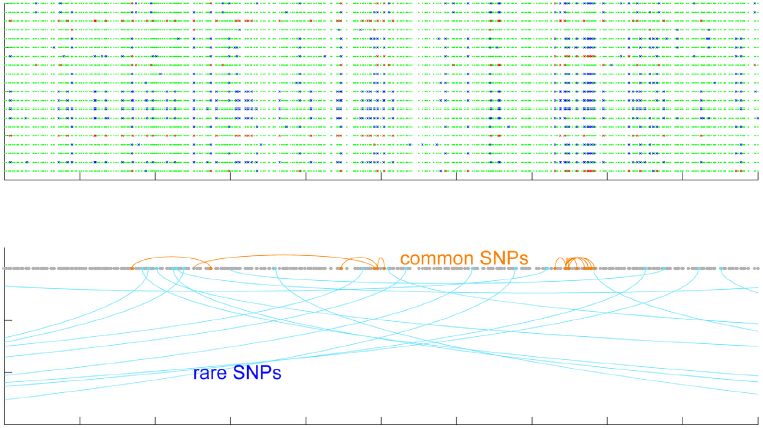
Genetic distance between a target SNP and its nearest neighbor. The top figure shows the genotypes of 20 samples in a 50-kb region. A green dot represents a homozygous reference site, blue is heterozygous, and red is homozygous alternate. We picked the SNPs with highest allele frequencies and lowest allele frequencies in this region, and marked them in orange and light blue in the bottom figure. The marked SNPs were connected with their nearest neighbor in terms of linkage disequilibrium. This figure clearly shows that common SNPs tend to have strong association with their nearby sites, while the sites associated with a rare SNP tend to be further away in genomic coordinates.

**Figure 8.**
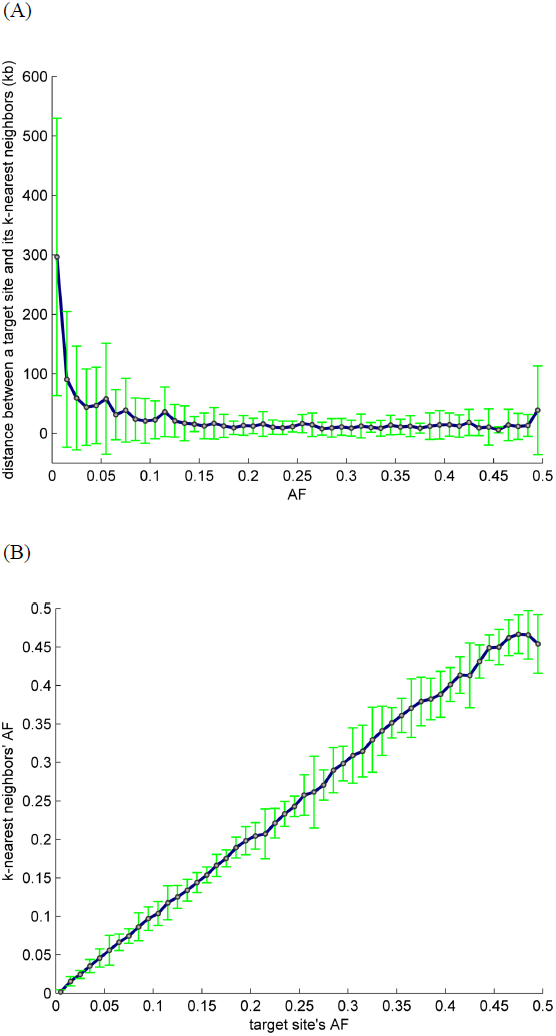
Motivation for the definition of nearest neighbors in Reveel. (A) Genetic distance between a target SNP and its k-nearest neighbors as a function of allele frequency. Rare SNPs exhibited very different behavior from common SNPs. (B) A strong correlation between the allele frequency of a target site and the allele frequencies of its k-nearest neighbors.

Hidden Markov model-based methods face a tradeoff between either explicitly modeling each rare haplotype, which leads to computational overhead due to the large number of parameters, or compressing the state space which leads to the loss of long-distance rare-haplotype LD information. In particular, previous state-of-the-art methods, such as MaCH and Thunder, apply a first-order Markovian model between two subsequent haplotypic positions. While such models have been demonstrated to work well for genotyping common variants, they face a challenge in modeling rare variants. On the lower side of the rare SNPs spectrum (⩽ 0.1%), the incorporation of LD information in previous methods did not improve the genotyping accuracy in the 1000 Genomes Project Phase 1; rather, the resulting genotyping accuracy was modestly lower than when not using LD information (The 1000 Genomes Project Consortium 2012). The explanation underlying this phenomenon may be as follows. Although rare haplotypes share common variants, they usually contain distinct rare variants that can serve as a signature. Leveraging those correlations within a simple Markovian model is impractical: every rare haplotype needs to be encoded in the model, captured as a distinct sequence of states in the HMM. The HMMs underlying currently available methods tend to eliminate rare alleles as noise, which contributes to biases towards homozygous reference. Conversely, to infer genotypes, our approach aims to identify the most informative sites in a way that is less sensitive to their genetic distance. The strategy is different from previous models that implicitly weaken the association between remote sites. By focusing on the most informative markers based on their LD, our method provides considerable improvement in the genotype calling of rare variants.

High AF SNPs are caused by one or more mutation events that occurred in the distant past; after many generations of recombination, the LD between high AF sites could become very complex. Therefore, perfect LD between a high AF site and a set of surrounding sites may not exist. In this particular case, genotype phasing on common variants is a useful complementary method to our genotype-calling algorithm. We incorporate a post-processing step into Reveel: after imputing the genotypes and genotype probabilities at likely polymorphic sites, we pick SNPs with AF > 1% and feed their genotype probabilities into Beagle (Browning and Browning 2009) for phasing. Finally the output dosages of Beagle are merged with the genotypes at rare SNPs.

An important feature of our algorithm at high AF sites is providing high-quality genotype probabilities. To demonstrate this point, we conducted an experiment for comparison, labeled as *Reveel-gatk-beagle*. In this experiment, we forced GATK to make calls across the sites identified by our algorithm with AF > 1%. Then, Beagle was trained on the outputs of GATK, producing dosages at these sites. Finally, we merged the outputs of Beagle and the genotypes called by our algorithm at rare SNPs for evaluation. The only difference between this approach and our default pipeline is which tool was used to created genotype probabilities. The comparison shown in Figure 9 clearly illustrates that Reveel outperforming *Reveel-gatk-beagle*.

**Figure 9.**
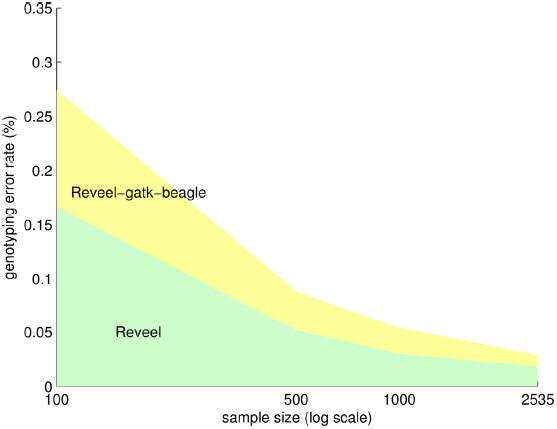
Impact of genotype probabilities produced by Reveel at high AF sites in estimate of Beagle’s output. For comparison, we forced GATK to make calls across the sites identified by Reveel with AF > 1% and to produce genotype probabilities at these sites. We replaced Beagle’s inputs in our pipeline with the genotype probability values obtained using GATK. The resulting pipeline was labelled as *reveel-gatk-beagle*.

The running time of Reveel scales linearly with the number of individuals *n* and linearly with the number of polymorphic sites *m* in our algorithms, expect that the process of estimating LD between every pair of polymorphic sites requires *m*^2^ computations. As we restrict the LD estimation within windows of size 500-kb – 1-Mb (see Methods), *m* is usually within the range of a few thousands depending on the size of the studied cohort, which results in practical running times in our experiments.

## Methods

The input to Reveel is a cohort of n sequenced individuals, for which read counts are available supporting each of the four possible nucleotides 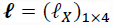 at every site in each sample. To genotype the individuals, Reveel performs the following four steps: (1) Polymorphic site discovery. A set of m putative polymorphic sites are identified across the genome. (2) Initialization of genotypes **G** = (*g*_*i,j*_) for every sample *i* and putative polymorphic site *j* identified in step 1. (3) Calculation of a matrix **P** = (*p*_*i,j,g*_), representing the probability of individual i having genotype *g* in position *j* given the current assignment **G**. (4) Calculation of new assignment **G**’ that maximizes the current entries of **P**; steps 3 and 4 are performed iteratively until convergence. (5) Final refinement of the genotypes **G**.

### Polymorphic site discovery

Knowing the set of observed reads supporting each of the four possible nucleotides 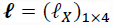 at a site in a sample, we can compute the probability that allele *X* ∈ {*A*, *C*, *G*, *T*} is present by marginalizing over possible genotypes given the read counts 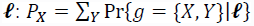, where the genotype *g* = {*X*, *Y*} is an unordered pair of alleles. The probability of genotype *g* given read counts can be computed as

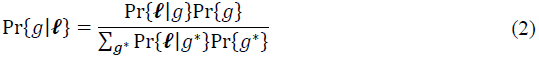

To compute the genotype probability, we first calculate the genotype likelihood, that is, the probability of observing 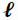 when the genotype is *g* = {*X*, *Y*}. The genotype can take one of ten possible assignments. The likelihood of a homozygous genotype can be written as a binomial probability mass function 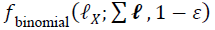, in which *ε* is the sequencing base error rate. The likelihood of a heterozygous genotype can be expressed as follows, in which the indicator function 1_condition_ equals to 1 if the condition is true; otherwise it equals to 0.

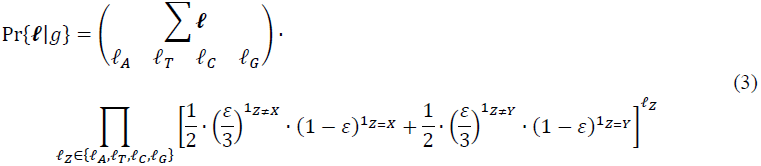

Assuming a non-informative prior over genotypes, the posterior 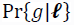 is proportional to the likelihood by Bayes’ rule.

We distinguish loci that contain true variations from those that arose from sequencing errors, as follows. Given a target locus and a candidate allele *X*, we define score_*X*_ representing the strength of the evidence for the existence of allele *X* at the target locus, using the summation of a monotonically increasing function over all the samples:

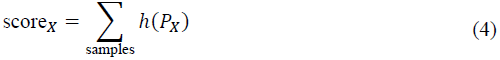

Assuming a site is monoallelic or bi-allelic, we define the allele with the highest score as the reference allele. The allele with the second highest score is a putative alternate allele. We distinguish alternate alleles from sequencing errors using a threshold score_th_.

The function *h* was trained using simulated annealing on simulated data sets in the design stage (different from the data sets used in the experiments), maximizing the overall recall under the perfect precision constraint. Since the function *h*(*z*) = *a* ∙ *z*/(1 + *a* − *z*) with a sole parameter *a* fit the training output well, this function was built in the tool and applied in all the experiments. We used the trained constant *a* = 5 × 10^−6^ in all our experiments.

### Genotype-calling algorithm

Given *m* candidate polymorphic sites that were identified in the previous step, we determine the genotypes of *n* samples simultaneously across the *m* sites. Let **G** be a *n* × *m* matrix, in which *g*_*i,j*_ = {0,1,2} represents the genotype of sample *i* at marker *j* being homozygous reference, heterozygous, homozygous alternate, respectively. Let **P** be a *n* × *m* × 3 matrix, where *p*_*i,j,h*_ represents the probability of *g*_*i,j*_ = *h*. We formulate the overall framework of our algorithm as a fixed-point model

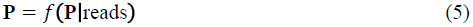

The function *f*(*z*) does not have a closed-form expression; instead, we estimate **P** by using a summarization-maximization iterative algorithm. This algorithm consists of two components: summarization and maximization. Given the genotype matrix **G**, in the summarization step, we estimate **P** in the context of LD and observed reads. In the maximization step, we update **G** with the genotypes associated with the largest probabilities within **P**. We iteratively apply these two components until convergence.

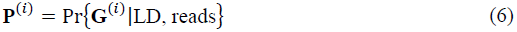

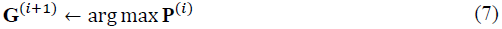

In each iteration we first apply the summarization on all markers and then apply the maximization on all markers. Using the subscript *target* to represent the marker being evaluated in a sample and 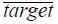 to represent all other makers in the same sample, we rewrite the above equations as:

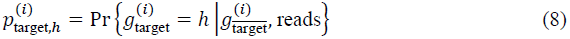

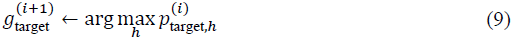

The main challenge is in the summarization step, where the LD information needs to be leveraged in a computational efficient way that also leads to high accuracy in estimating the conditional probabilities. Here, we introduce a technique that leverages the most informative markers in terms of LD. For each marker, we find its *k*-nearest neighbor markers (as defined in the next section) in terms of LD. Equation 8 is replaced by:

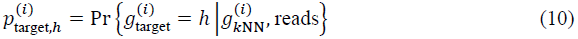

The observed reads provide two forms of evidence: the read counts supporting alleles at the target marker (denoted as *r*_target_) and the allele frequencies at the evaluated marker across samples (denoted as *θ*). To utilize the read counts, by the chain rule, we rewrite the conditional probability in Equation 10 to yield Equation 11.

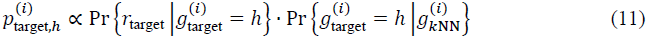

The calculation of the first term is straightforward. To calculate the second term, we use the probability of genotypes in the *i*-th iteration as follows. For each sample *j*, we calculate the probability that this sample has genotype *h* at the target locus and genotypes 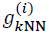 at the neighbor loci. We use subscript (target, *j*) to represent the marker on the same locus as the target but in sample *j*, distinguished from the target SNP being evaluated. Similarly, we use subscript (*k*NN, *j*) to represent the *k*-nearest neighbors in sample *j*. With these notations, the above probability can be expressed 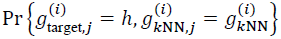. Summing this probability over all the samples yields the expected sample count *C*_*h*_ with genotype *h* at the target SNP and 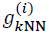 at the neighbors; summing over all the samples and all the possible *h*’s yields the expected count *C* having 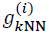 the neighbors. We use the ratio *C*_*h*_/*C* as the new conditional probability 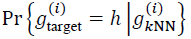.

In practice, because the sample size is usually limited to hundreds or thousands, the conditional probability assessment could be biased (Friedman et al. 1997), which can significantly affect the performance. To reduce bias, we use Laplace smoothing (Hansen et al. 2005). In summary, the second term is given by the following expression, in which we set *t* = 1 if AF ⩽ 1% and *t* = 0.01 otherwise.

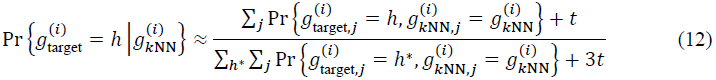

Although Laplace smoothing is used, if the initial **G**^(0)^ is biased towards the homozygous reference on certain markers, then 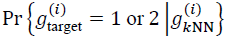 tends to be a very small number. Thus, the converged results are also very likely to be biased. To address this issue we leverage the other signal given by the reads, that is, the alternate allele frequency over samples, and rewrite Equation 11 as:

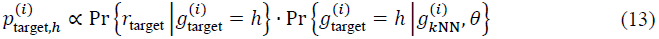

Once again, we face the problem of assessing the conditional probability in the second term,but this time we obtain knowledge from an additional source. Let 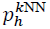 be 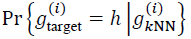, and let 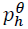 be 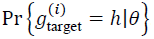. The probabilities evaluated from different sources are combined using a noisy-MAX gate (Zagorecki and Druzdzel 2013). The expressions are as follows.

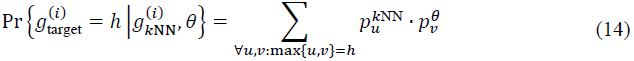

In contrast to the estimate provided by Equation 12, which is biased towards homozygous reference, this estimate is biased towards homozygous alternate. We use each of the above two estimates alternatingly in the iterations of our summarization-maximization algorithm.

### Nearest neighbor calculation

To define the *k* nearest neighbors of a locus, we introduce three metrics to approximate the LD between two loci. As this evaluation is performed on every pair of candidate polymorphic sites, we need metrics with low computational overhead. Commonly used metrics such as the correlation coefficient require the estimation of genotypes based on the observed reads; this estimation requires a considerable computational cost. The main benefit of the metrics we present here is that they can be directly applied on the read counts.

Let *S*_*i*_ be a set of samples that have at least one read at locus *i* supporting alternate alleles. The first metric is defined as the Jaccard index of two sets

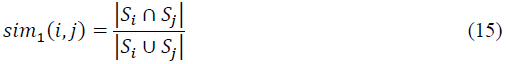

This metric utilizes the presence of reads that support alternate alleles. As a second, more informative metric, we apply the Jaccard index on multisets, accounting for repeated elements. Set 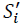 is defined as the collection of *r*_*i,t*_ copies of *t*’s, where *r*_*i,t*_ is the number of reads at locus *i* of sample *t* supporting alternate alleles. The second metric is thus

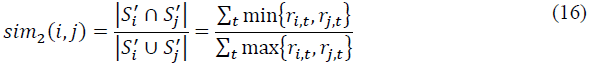

Finally, we define the third metric that produces a more rapidly increasing score as both samples exhibit more reads that support alternate alleles:

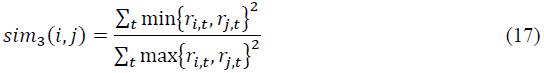

We apply the summarization-maximization algorithm separately, leading to matrices **P**_*i*_ for *i* = 1, 2, and 3. Then, we combine the three matrices by using the average probability at each marker (as called as the mean combination rule) (Kittler et al. 1998, Xu et al. 1992):

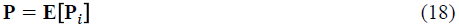

The combined genotype matrix is given by

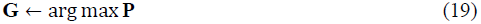

The inter-marker LD at most extends to a few hundred kilobases (kb) (Reich et al. 2001, Schaffner et al. 2005). To compute nearest neighbors efficiently, we tile the genome with a set of non-overlapping blocks. The *k*-nearest neighbor markers are selected from the block to which the target marker belongs. We found that block sizes of 500-kb to 1-Mb results in a good balance between accuracy and running time. Our default block size is 1-Mb.

The impact of parameter *k* on the quality of the approximation in Equation 12 is significant. Let 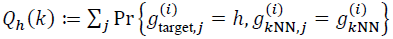 and 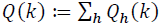. An overly large *k* can result in very small *Q*(*k*) and therefore low-quality conditional probability tables. Assuming LD, *Q*(*k*) can be very roughly estimated as *n* ∙ [(1 -*maf*)^2^]^*A*^ ∙ [2 ∙ *maf* ∙ (1 -*maf*)]^*B*^ ∙ [*maf*^2^]^*C*^, where *A*, *B*, *C* are the counts of 0, 1, 2 in the genotype pattern 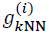 and *A* + *B* + *C* = *k*. In other words, given a fixed sample size *n*, *Q*(*k*) exponentially shrinks with the increment of *k*. Based on our experiments, we recommend the following settings: *n* ⩽ 75, *k* = 2; 75 < *n* ⩽ 250, *k* = 3; *n* > 250, *k* = 4. As cohorts become considerably larger than 1KGP in the future, we expect that larger values for *k* will yield better performance.

Finally, the conditional probability computed with different values for *k* conveys LD on different levels. To balance the impact of the selection of *k*, we rewrite Equation 11 as

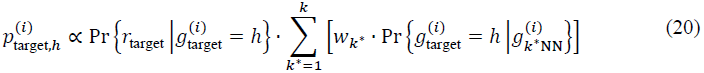

where the weight *w*_k^∗^_ can be 1/*k*^∗^ or 2*k*^∗^/(*k* + *k*^2^). In our experiments, we use Equation 20 with *w*_k^∗^_ = 1/*k*^∗^.

### Initial genotypes

Given a low-coverage sequencing data set, we observe only a few (if any) reads at a target site. Thus, using these reads to estimate 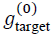 is not a good initial guess. Instead, we use the reads at the *k* nearest loci to amplify the low-coverage data. More formally, let *r*_*i*_ and 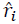be the number of reads at locus *i* supporting alternate and reference alleles. Instead of using *r*_target_ and 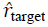, we use 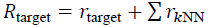 and 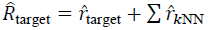 for the initial guess, which is equivalent to amplifying the depth of the target site. We assign

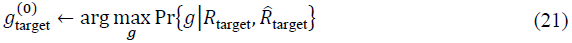

### Final refinement

The method described in the previous sections achieves sufficiently high performance with a very limited number of neighbor SNPs. To further improve the genotyping accuracy, we apply a phasing method to the common and low frequency SNPs (AF ⩽ 1%). Since previous publications have proposed high-quality phasing algorithms (Browning and Browning 2009, DePristo et al. 2011, McKenna et al. 2010), we use BEAGLE (Browning and Browning 2009) for this step. We feed the genotype likelihoods of high-frequency SNPs into BEAGLE, and then merge the phased dosage into our outputs.

